# A comprehensive influenza reporter virus panel for high-throughput deep profiling of neutralizing antibodies

**DOI:** 10.1101/2020.02.24.963611

**Authors:** Adrian Creanga, Rebecca A. Gillespie, Brian E. Fisher, Sarah F. Andrews, Liam Hatch, Tyler Stephens, Yaroslav Tsybovsky, Michelle C. Crank, Adrian B. McDermott, John R. Mascola, Barney S. Graham, Masaru Kanekiyo

## Abstract

A number of broadly neutralizing antibodies (bnAbs) to influenza virus have been isolated, characterized and developed as potential countermeasures for seasonal influenza epidemic and pandemic. Deep characterization of these bnAbs and polyclonal sera is critical to our understanding of influenza immunity and for desgining universal influenza vaccines. However, conventional influenza virus neutralization assays with live viruses require high-containment laboratories and are difficult to standardize and roboticize. Here, we built a panel of engineered influenza viruses carrying a fluorescent reporter gene to replace an essential viral gene. This restricts virus replication to cells expressing the missing viral gene *in trans*, allowing it to be manipulated in a biosafety level 2 environment. Using this system, we characterize the neutralization profile of a set of published and new bnAbs with a panel consisting of 55 viruses that spans the near complete antigenic evolution of human H1N1 and H3N2 viruses, as well as pandemic viruses such as H5N1 and H7N9. Our system opens opportunities to systematically characterize influenza immunity in greater depth, including the response directed at the viral hemagglutinin stem, a major target of universal influenza vaccines.

## Introduction

Influenza virus continues to cause seasonal epidemics and pandemics despite vaccine availability. Between 2010 and 2018, seasonal influenza virus infection resulted in an estimated 140,000–810,000 hospitalizations and 12,000–61,000 deaths in the United States alone(*1*). In the most recent influenza pandemic in 2009, the virus claimed >10,000 lives(*2*). In addition, highly pathogenic avian influenza viruses cause sporadic outbreaks in humans with a high mortality rate, posing potential risk of human adaptation and are considered pandemic threats(*3*). Of the four types of influenza viruses (A–D), only influenza A and B viruses cause high morbidity and mortality in humans. Hemagglutinin (HA) and neuraminidase (NA) are the only targets for neutralizing antibodies on the virus envelope. HA is required for viral entry into the host cell through binding to sialic acid moieties on glycoproteins or glycolipids and mediates fusion of the viral membrane with host endosomal membranes. NA cleaves sialic acid and promotes the release of progeny viruses from infected cells. Neutralizing antibodies are primarily directed against HA and can either compete for receptor-binding, inhibit the membrane fusion machinery, or cause virus aggregation. Antibodies to NA can sometimes have neutralizing activity, but are thought to primarily limit virus egress by inhibiting enzyme activity and release of virus thereby inhibiting viral spread. Based on the genetic and antigenic properties of HA and NA, influenza A viruses are divided into group 1 and 2, each of which have several subtypes defined primarily by HA. To date, there are 18 HA and 11 NA subtypes identified and characterized(*4*). Currently, strains of H1N1 and H3N2 influenza A viruses co-circulate in humans. In addition, several other subtypes of animal influenza viruses (e.g., H5N1, H6N1, H7N9, H9N2, and H10N8) can also infect humans and occasionally result in mortality. In contrast to influenza A viruses, which have an avian reservoir and can achieve sustained transmission in several mammalian species (e.g., swine, dogs, horses, and bats), influenza B viruses are isolated almost exclusively from humans with a more limited evolutionary history and have diverged into only two genetically and antigenically-distinct lineages (Victoria- and Yamagata lineages), which presently co-circulate in humans(*5*).

Due to the wide diversity and continuous evolution of influenza viruses, repeated vaccinations remain the most effective approach to mitigate influenza burden in humans. Current influenza vaccines confer protection primarily to the strains closely related to those used in its preparation, thus reformulation is needed each season to match the antigenic properties of circulating strains. As a result, the effectiveness of seasonal influenza vaccines varies widely (<10–60%) depending on the adequacy of this match(*6, 7*). Moreover, current seasonal influenza vaccines confer no protection against animal viruses that can cause human diseases, such as H5N1 or H7N9 highly pathogenic avian influenza viruses in animal models. There are also limitations in current manufacturing approaches and general public apathy for immunization. Therefore, new vaccine options for influenza are urgently needed and have become a high priority for public health officials(*8*). The current approach for qualifying vaccines each year and licensing vaccines for public use is largely based on the ability of new or reformulated vaccines to induce neutralizing antibodies targeting the receptor-binding site (RBS) in the HA head, which have hemagglutination inhibition (HAI) activity. The HAI assay depends on the availability of HA to bind receptors on red blood cells and was developed as a surrogate for measuring neutralizing activity in the 1940s. The HAI assay is also complicated because some influenza strains do not have enough hemagglutination activity to be useful in this assay.

The discovery of broadly neutralizing antibodies (bnAbs) capable of neutralizing multiple influenza virus subtypes in humans opens an opportunity for developing a universal influenza vaccine which elicits such antibodies(*9–11*). Many of these antibodies target conserved epitopes in the HA stem and neutralize virus by inhibiting the viral fusion machinery and hence, the activity is not detectable by HAI assay. Several less broad bnAbs bind to the RBS in the HA head and prevent the virus from engaging sialic acid moieties on the host cell surface, thus exhibiting HAI activity(*12*). Currently, most universal influenza vaccine candidates aim to induce protective levels of such bnAb response(*11, 13*). Therefore, comprehensive analysis of the neutralization breadth and potency regardless of HAI activity is critical to accelerate the efforts to develop effective universal influenza vaccines.

The influenza microneutralization (MN) assay is the most commonly used assay to measure virus neutralizing activity of antibodies to influenza virus(*14*). The assay measures virus replication by detecting viral nucleoprotein (NP) levels with an enzyme-linked immunosorbent assay (ELISA), titrating hemagglutination titers, or through monitoring virus-induced cytopathic effects. Plaque-reduction neutralization assay is also commonly used for influenza and other viruses. These approaches are labor-intensive, not easily scalable and the handling of live viruses of animal origin requires high-containment laboratories. There is also significant performance variability between laboratories due to multi-step signal amplification or reliance on experienced operators. Given these inevitable limitations of the current MN assay, there is a need to transform the MN assay to be safe, high-throughput, robust, easy to standardize, and automation compatible(*15*).

The use of reporter viruses for developing high throughput and reproducible neutralization assays has greatly advanced our ability to measure and characterize antibody responses induced by infection and/or vaccination(*16–18*). While pseudotyped reporter lentiviruses have been constructed for influenza and utilized effectively for rank-ordering neutralizing activity (*19*), they are often criticized for being too sensitive and may not accurately represent some of the key properties of influenza viruses like entry through the endosome under low pH conditions(*20, 21*). Therefore, building influenza viruses with a reporter feature is an attractive alternative. Amongst several approaches to produce influenza reporter viruses, a replication-competent reporter virus can be developed by fusing a reporter gene to a viral gene (e.g. PA or NS)(*22*). This new virus remains as virulent and replication-competent as the parental virus, making it suitable for viral pathogenesis studies, yet still subject to precautions regarding handling in high-containment laboratories. In contrast, single-cycle infectious or replication-restricted reporter (R3) influenza viruses can be prepared by replacing one of the essential viral genes (e.g. PB1, or HA) with a reporter gene(*22–25*). Thereby the R3 virus is capable of replicating only in cells engineered to express the deleted viral gene *in trans*, making it safe to rescue and propagate in low-containment laboratories.

In the present study, we developed a comprehensive panel of R3 viruses to enable high-throughput and in-depth influenza virus neutralization profiling. We generated a robust neutralization matrix of 24 monoclonal antibodies and 55 R3 viruses spanning 6 subtypes of influenza virus, and compared the R3 virus-based assay with the gold-standard ELISA-based MN assay(*14*). The reporter virus assay provides a more robust method for probing the breadth of anti-influenza immunity needed for developing universal influenza vaccines.

## Results

### Generation of R3ΔPB1 viruses

The influenza virus has a segmented genome, which allows its rapid and relatively easy genetic manipulation. Each segment consists of either one or two protein coding sequences flanked by short noncoding regions (NCRs). The genome packaging signal sequences are located at both 3’- and 5’-termini of each segment encompassing the entire NCRs and part of the adjacent protein coding sequence. Genome packaging sequences are unique to each segment and required for an efficient incorporation of each viral RNA molecule into the virion. The PB1 segment encodes two proteins: PB1, the main component of the viral RNA-dependent RNA polymerase, and PB1-F2, translated from the +2 frame and not required for virus replication(*26*). The protein coding sequence of PB1 (A/WSN/1933, H1N1) has 2,274 nucleotides and is flanked by 24 nucleotides at the 3’- and 43 bases at 5’-termini. In addition to the NCRs, genome packaging signals of the PB1 segment comprise 120 bases of the coding region at both 3’- and 5’-termini(*23, 27*) (Figure S1).

To build R3 influenza viruses, we altered the PB1 segment by removing the PB1 coding sequence not required for genome packaging and replacing it with the reporter encoding fluorescent protein tdKatushka2(*28*). To prevent translation of alternative transcripts from PB1 segments, we mutated potential initiation codons (ATGs) found between the 3’ end and the reporter open reading frame (ORF). PB1 protein is essential for virus replication therefore, R3 influenza viruses in which PB1 protein was removed (R3ΔPB1) can be propagated only in cells expressing the PB1 *in trans* (Figure 1B). Thus, we used MDCK-SIAT1 cells, which constitutively express human β-galactoside α2,6-sialyltransferase 1 (SIAT1)(*29*) to prepare a cell line stably expressing PB1 of A/WSN/1933 by transfecting a plasmid encoding a puromycin-resistance gene, PB1, and a self-cleaving peptide derived from Thosea asigna virus 2A (T2A) in between the two genes. The R3ΔPB1 virus was rescued by reverse-genetics using 8 plasmids encoding HA and NA of wild-type H1N1 or H3N2 influenza A viruses; PB2, PA, NP, M and NS of A/WSN/1933; and another encoding the reporter. To validate the use of R3ΔPB1 viruses for influenza neutralization assays, we also rescued replication-competent H1N1 and H3N2 parental viruses, which possess HA and NA of wild-type viruses and the internal genes including PB1 of A/WSN/1933, and were propagated in MDCK-SIAT1 cells (Figure 1B).

**Fig. 1.**
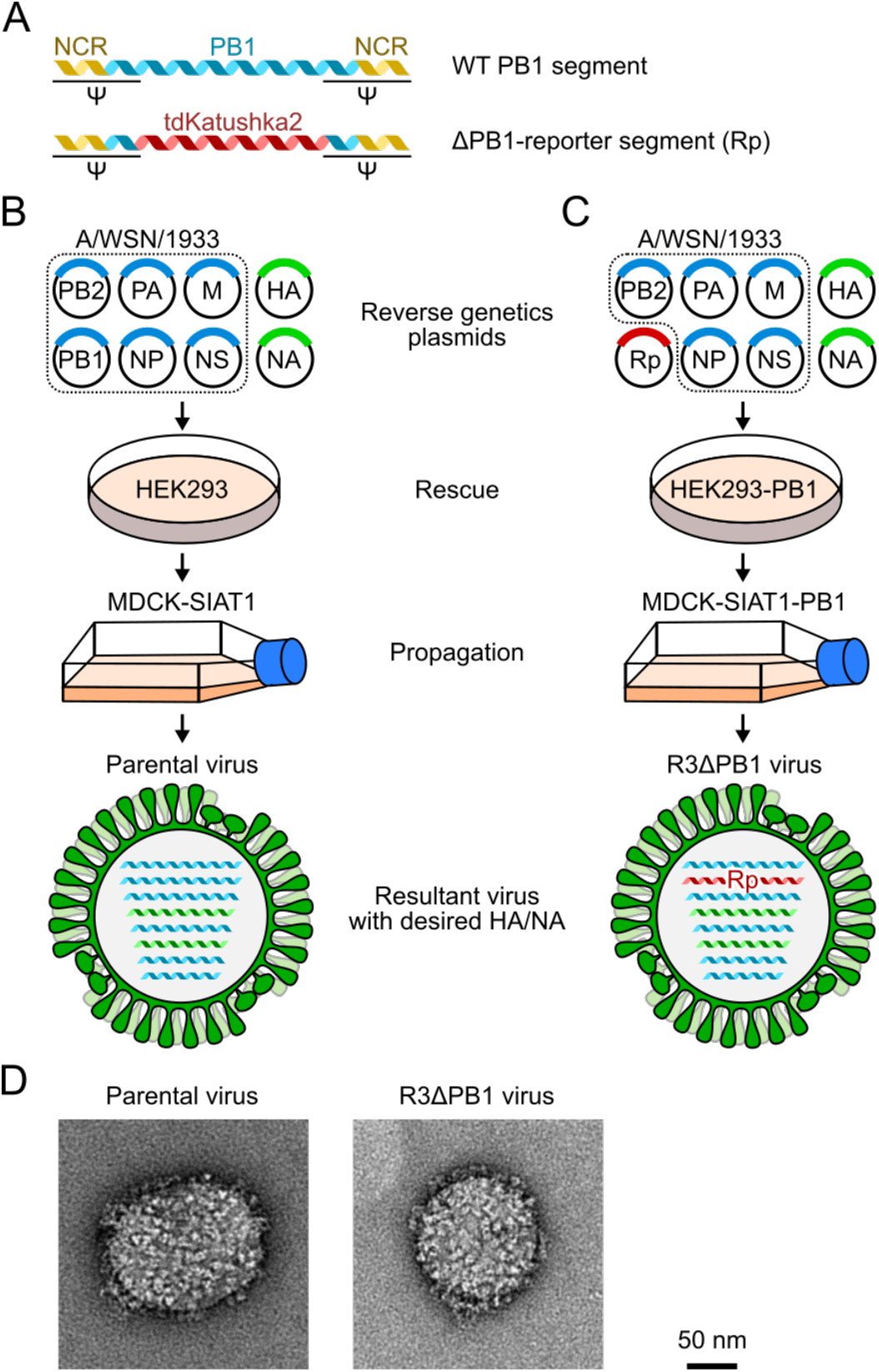
Development of replication-restricted reporter (R3) ΔPB1 influenza A virus. (A) Design of engineered PB1 segment encoding fluorescence reporter tdKatushka2. Rescue of parental wild-type molecular clone (B) and R3ΔPB1 (C) viruses using reverse genetics of eight plasmid system with HA and NA segments of virus of interest and internal gene segments of A/WSN/1933. PB1 segment of A/WSN/1933 is used to rescue molecularly cloned parental viruses, while engineered PB1 segment is used for R3ΔPB1 influenza viruses. R3ΔPB1 influenza viruses require cells expressing PB1 *in trans* for rescue and propagation. (D) Negative stain electron microscopy of parental molecular clone (left) and R3ΔPB1 (right) of A/Michigan/45/2015 (H1N1) virus.

To examine the morphology of R3ΔPB1 virus (A/Michigan/45/2015, H1N1), we performed negative stain electron microscopy and found that there was no visible difference in size and spike density between R3ΔPB1 and the corresponding wild-type molecularly cloned parental viruses, indicating that the R3ΔPB1 virus remains morphologically indistinguishable from its parental virus (Figure 1D). Virus growth kinetics of the R3ΔPB1 (A/Michigan/45/2015) and its parental viruses in MDCK SIAT1 and PB1-expressing MDCK-SIAT1 cells confirmed that the R3ΔPB1 virus replicates only in PB1-expressing cells, whereas parental virus replicates similarly in MDCK-SIAT1 cells with or without PB1 expression (Figure S2). When we compared virus infection rate between the two viruses over time we detected approximately the same number of infected cells (cca. 5%) at 18 h post infection. These results demonstrate that the R3ΔPB1 viruses cannot replicate in cells lacking PB1 expression and replicate comparably in PB1-expressing cells to corresponding parental viruses carrying PB1 with identical HA and NA.

### Influenza virus neutralization assay using R3 virus

We tested whether the R3ΔPB1 viruses can be used in an influenza virus neutralization assay. To do so, neutralization assays were carried out utilizing both R3ΔPB1 and parental viruses with matched HA and NA, where we compared the inhibitory concentration of a set of monoclonal antibodies (mAbs). To verify the reporter-based readout, the R3ΔPB1 virus-infected cells were detected either by ELISA with anti-NP antibody or by fluorescence using an automated image-based plate reader at 24 h post infection. We measured neutralizing activity (80% inhibitory concentration, IC_80_) detected by both ELISA- and fluorescent reporter-based readouts for several mAbs against a total of eight R3ΔPB1 viruses (4 H1N1 and 4 H3N2 viruses) and found a strong positve correlation (Pearson r = 0.94, p < 0.001) between the two IC_80_ datasets, demonstrating that the fluorescent reporter-based readout yields comparable neutralization results to conventional ELISA-based detection method (Figure 2A and S3).

**Fig. 2.**
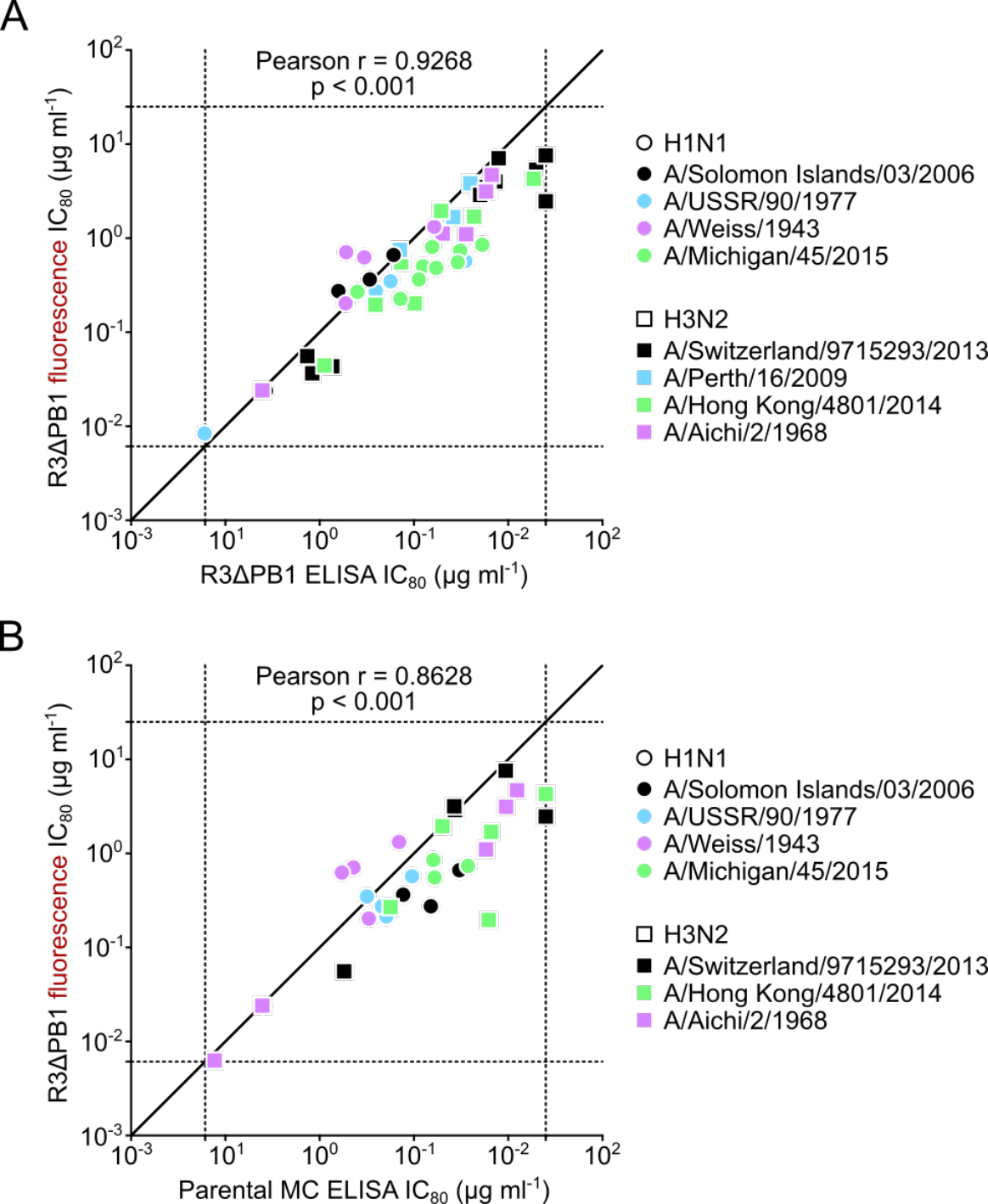
Assement of R3ΔPB1 virus for neutralization assay. (A) Correlation between fluorescence- and ELISA-based readouts. Each dot indicates the neutralization titers (IC_80_ μg/ml) of a single monoclonal antibody against an R3ΔPB1 influenza virus measured by fluorescence readout (y-axis) and ELISA with anti-NP antibody (x-axis). (B) Correlation between assays using R3ΔPB1 and molecular clone viruses. Each dot indicates the neutralizing activity (IC_80_ μg/ml) of a single monoclonal antibody using R3ΔPB1 (y-axis) and parental viruses (x-axis) expressing the same HA and NA.

To assess the R3ΔPB1 viruses with the MN assay format, we next determined neutralizing IC_80_ for 6 mAbs against matched pairs of R3ΔPB1 and parental viruses (4 H1N1 and 3 H3N2 viruses). Cells infected with R3ΔPB1 viruses were detected by fluorescence, while cells infected with parental viruses were detected by ELISA. There was a positive correlation (Pearson r = 0.87, p < 0.001) between neutralization IC_80_ of mAbs against R3ΔPB1 and parental viruses, showing that R3ΔPB1 viruses retained the neutralization sensitivity of parental viruses with matched HA and NA (Figure 2B). In conclusion, our studies indicate the neutralization assay using R3ΔPB1 viruses with fluorescence-based readout can be used as high-throughput, safe, and reliable measurement of virus neutralizing activity.

### Building a comprehensive panel of R3 viruses

We aimed to build a comprehensive panel of R3ΔPB1 viruses spanning the entire antigenic evolution of human H1N1 and H3N2 subtype viruses. We selected representative influenza strains based on phylogenetic analysis of HA sequences deposited in public databases, literatures on genetic and antigenic evolution of human H1N1 and H3N2 influenza viruses, and vaccine strains utilized since 1930s(*28, 30–33*).

H1N1 subtype virus was introduced into the human population in 1918 and circulated until it was replaced by H2N2 virus in 1957. H1N1 virus reemerged in 1977 and circulated until 2009. During this period, H1N1 viruses evolved significantly through genetic drift into multiple clades with distinct genetic and antigenic properties(*32, 34–36*). To capture the antigenic variations of these viruses we chose 7 matched HA and NA sequences from viruses circulating between 1933 and 1957, and 13 HA and NA sequences from viruses circulating between 1977 and 2009. The 2009 pandemic H1N1 virus acquired sustained human-to-human transmission and rapidly and completely replaced the pre-pandemic H1N1 virus. Since its emergence, the 2009 pandemic H1N1 virus has accumulated several amino acid substitutions linked to changes in antigenicity(*37*). Therefore, we included 3 HA and NA sequences from viruses isolated between 2009 and 2015, and HA and NA sequences of swine-origin A/New Jersey/8/1976 (H1N1), which caused an isolated outbreak in 1976 in the United States. In summary, our H1N1 panel consists of 20 pre-pandemic strains, 3 pandemic strains, and one swine-origin H1N1 strain (Figure 3).

**Fig. 3.**
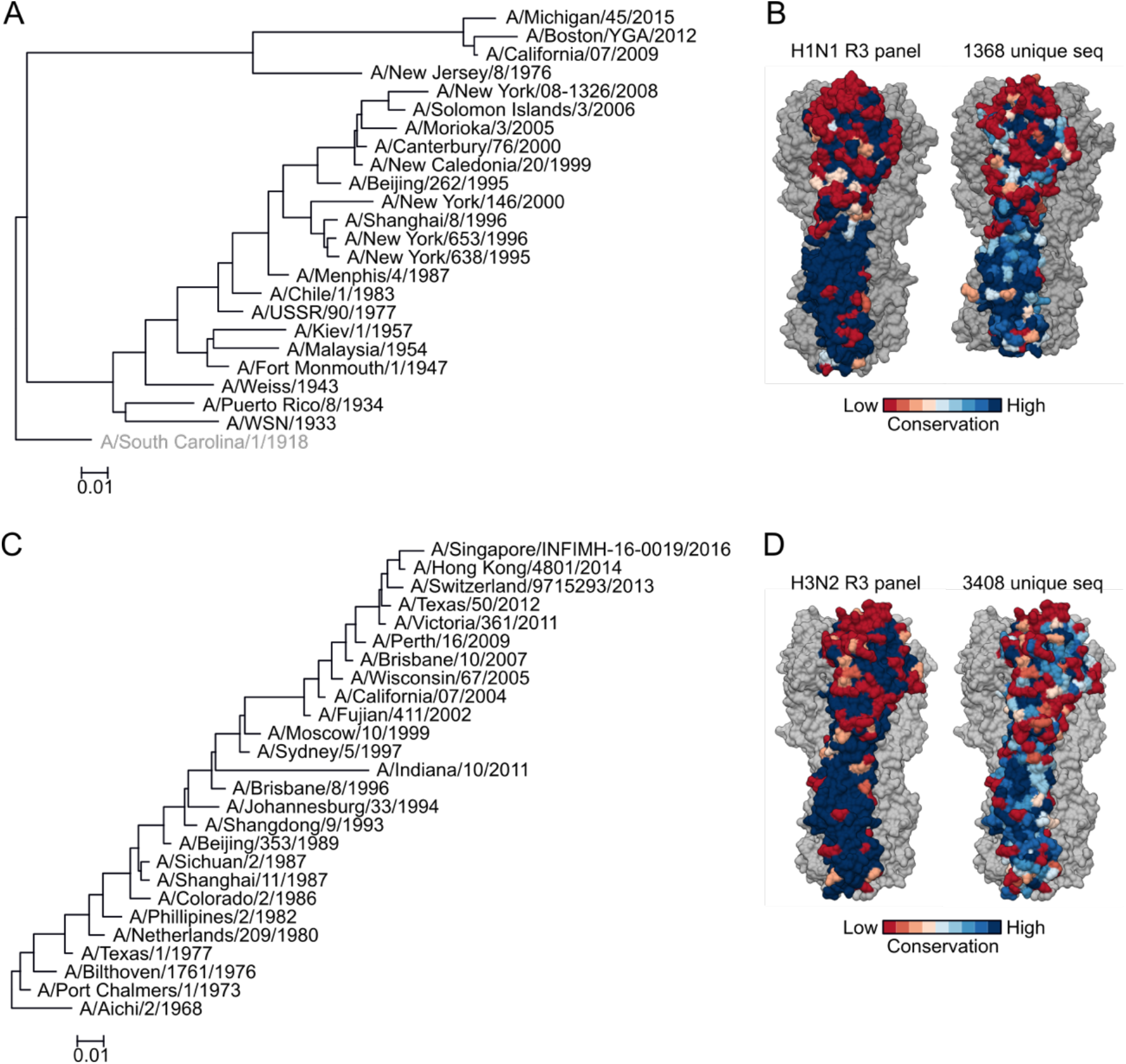
Phylogenetic and protein surface conservation analysis of HA of H1N1 and H3N2 viruses used in the study. Phylogenetic tree of 24 H1 HA sequences used in the R3 panel (A). The tree was generated by NJ method and rooted with A/South Carolina/1/1918 (not used in the study). (B) Protein surface conservation of H1 HA used in the R3 panel and a larger non-redundant representative dataset. Phylogenetic tree of 26 H3 HA sequences used in the R3 panel. The tree was rooted with A/Aichi/2/1968. (D) Protein surface conservation of H3 HA used in the R3 panel and a larger non-redundant representative dataset. Amino acid conservation at each residue was calculated and scored by using the Consurf server (https://consurf.tau.ac.il/) and rendered on the structure of A/California/07/2009 (PDB: 3LZG) (B) and A/Victoria/361/2011 (PDB:4WE8) (D).

H3N2 subtype virus has been circulating in humans since its emergence in 1968. Comprehensive genetic and antigenic analysis of human H3N2 viruses groups them into 14 distinct antigenic clusters(*26, 31, 33, 38*), and hence, we include several representatives from each antigenic cluster in our panel. Sporadic outbreaks with swine H3N2 variant (H3N2v) have also been described in humans(*39*). Therefore, we included the swine-origin H3N2v strain, A/Indiana/10/2011. As a result, our H3N2 panel includes 25 human H3N2 strains and one H3N2v strain (Figure 3).

When we calculate the conservation of solvent-exposed surface amongst H1 and H3 HAs included in our panel, we notice that the head region of HA of both H1 and H3 is substantially more variable while the stem region is mostly conserved as expected (Figure 3B,D). The degree of surface conservation reflects that of much larger datasets of H1 and H3 HAs (Figure 3B,D)(*40*), suggesting that the selected HA sequences in our R3 virus panel capture the HA diversity of human H1N1 and H3N2 viruses.

### Generation of R3ΔHA viruses with highly pathogenic influenza virus sequences

Working with highly pathogenic influenza viruses (e.g., H5N1, H7N9, 1918 H1N1) or influenza lineages disappeared from human population (e.g., H2N2) requires high containment laboratories. Although R3ΔPB1 viruses have limited capacity to replicate due to the requirement of PB1 complementation, they retain the ability to reassort the HA segment with wild-type influenza viruses. To prevent this event, we evaluated alternative, approaches less susceptible to reassortment and generated reporter viruses expressing HA and NA of viruses with pandemic potential(*24, 41*).

First, we developed influenza viruses, unable to reassort HA segments, by making “rewired” replication-restricted reporter ΔPB1 viruses (R4ΔPB1) as previously described(*41*). For this purpose, two segments of influenza genome were altered: the PB1 segment was modified to encode HA, and the HA segment was modified to encode the tdKatushka2 reporter. Using these altered PB1 and HA segments and reverse genetics, we were able to rescue R4ΔPB1 A/Switzerland/9715293/2013 (H3N2) virus in PB1-expressing cells (Figure S4A–C). By measuring neutralizing IC_80_ of 15 mAbs for both R3ΔPB1 and R4ΔPB1, we found a strong positive correlation (Pearson r = 0.94, p < 0.001) between IC_80_ neutralizing activity against the two viruses, demonstrating that these viruses have equivalent neutralization sensitivities (Figure S4D). Moreover, we found that the R4ΔPB1 A/Switzerland/9715293/2013 (H3N2) virus could not reassort its HA segment with A/Solomon Islands/03/2006 (H1N1) when the two viruses were co-infected and propagated on PB1-expressing cells (Figure S4E). Although these results show that R4ΔPB1 viruses can be safely rescued and used for high-throughput neutralization assay, further work is required to optimize the rescue and the propagation of R4ΔPB1 to be a practical and deployable process for producing reporter viruses.

Next, we explored an alternative approach to rescue R3 viruses unable to reassort the HA segment by replacing the HA coding sequence with the reporter gene (R3ΔHA). In this configuration, the virus lacks a functional HA segment and is unable to reassort the segment with other viruses (Figure 4A–C), and it will be able to replicate only in HA-expressing cells(*24*). To assess R3ΔHA viruses for the reporter neutralization assay, we generated R3ΔHA A/Switzerland/9715293/2013 (H3N2) virus was generated to compare the neutralization IC_80_ of several mAbs against both R3ΔPB1 and R3ΔHA viruses. Thre was a positive correlation between neutralization IC_80_ determined with these viruses (Pearson r = 0.90, p < 0.001), although we noticed slight differences in sensitivity to neutralization when testing anti-stem antibodies against R3ΔPB1 and R3ΔHA viruses (Figure 4D). Using this approach, we prepared 6 R3ΔHA viruses (i.e., 2 H5N1, 2 H7N9, 1 H2N2, and 1 H10N8) in addition to 49 R3ΔPB1 virus panel.

**Fig. 4.**
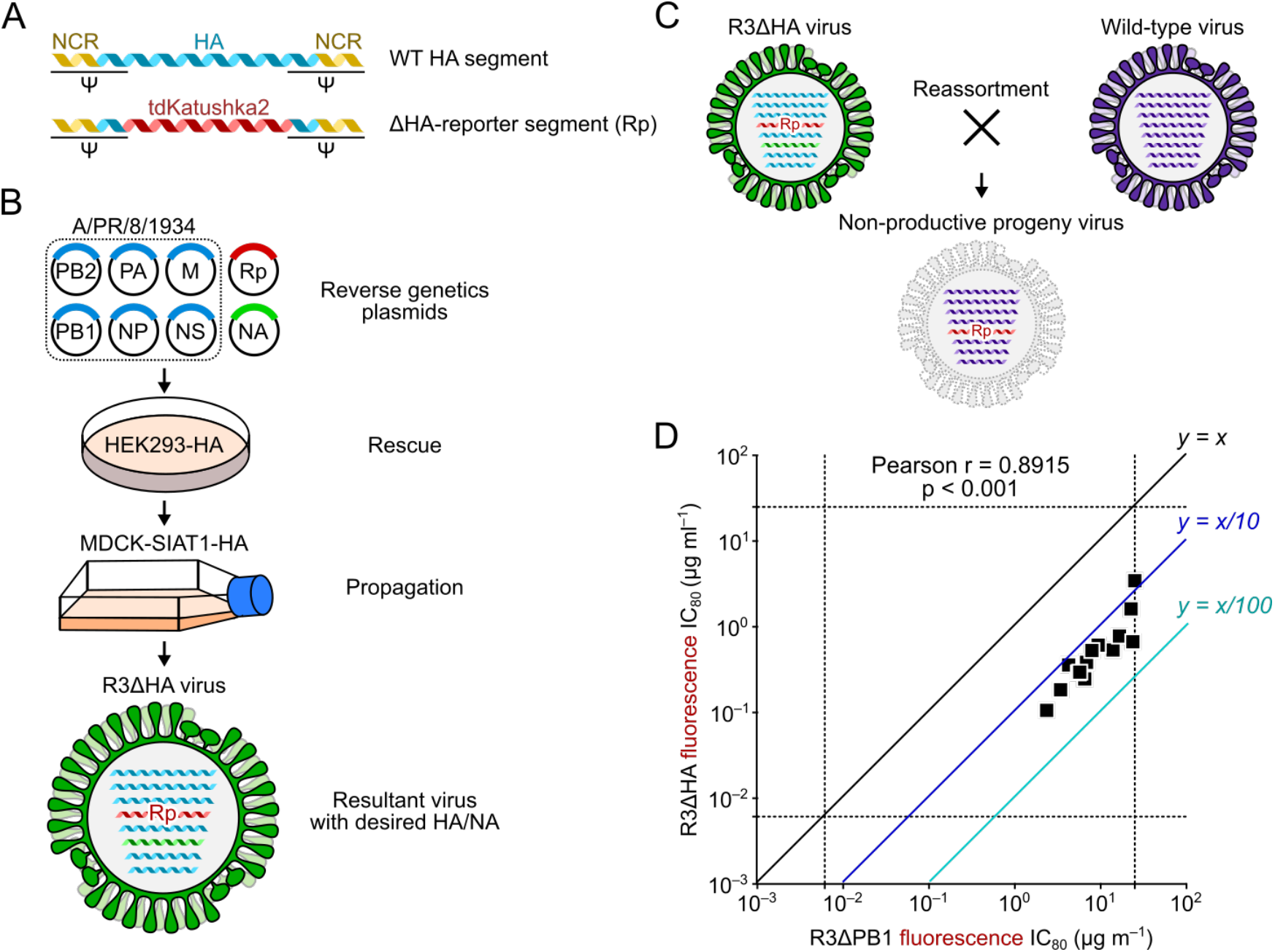
Development of R3ΔHA influenza viruses. (A) Design of HA segment used to rescue R3ΔHA virus. (B) R3ΔHA virus requires cells expressing HA protein *in trans* for rescue and propagation. (C) Non-viable reassortment between R3ΔHA and wild-type influenza viruses. Reassortant virus carrying the engineered HA segment encoding the reporter gene results in replication-deficient virus. (D) Neutralization sensitivity of R3ΔHA viruses. Correlation between neutralization titers of 24 monoclonal antibodies against R3ΔPB1 and R3ΔHA of A/Switzerland/9715293/2013 H3N2 is shown. Each dot indicates titers (IC_80_ μg/ml) of a single monoclonal antibody against R3ΔHA (y-axis) and matched R3ΔPB1 viruses (x-axis).

### Profiling of influenza neutralizing monoclonal antibodies with a 55-virus panel

Using our comprehensive panel of R3 influenza viruses spanning 6 different influenza A subtypes, we profiled the neutralization breadth and potency of a total of 24 human mAbs. Thirteen of them were isolated at the Vaccine Research Center from peripheral blood mononuclear cells collected as part of the H5N1 or H7N9 vaccine clinical trials(*40, 42–44*), while 11 other antibodies were previously described elsewhere(*45–54*). Amongst 24 antibodies, 17 antibodies recognize epitopes on the conserved HA stem region and 7 antibodies (i.e., CH65, 5J8, C05, F045-092, F005-126, 315-19-1D12, and 310-33-1F04) bind within the HA head region. We generated a matrix of neutralizing profiles for 24 antibodies against 55 R3 viruses consisting of 1,320 data points and analyzed the matrix by hierarchical clustering (Figure 5). Despite the fact that there is no associated information about viruses in the input dataset, the profile matrix segregates not only groups of viruses but also virus subtypes into defined clusters (Figure 5). Interestingly, the hierarchical clustering groups several antibodies according to their immunogenetic composition or convergent antibody class(*44, 55, 56*). For example, MEDI8852 is clustered with 2 other antibodies (315-53-1F12 and 315-53-1B06) and all the three antibodies belong to the V_H_6-1+D_H_3-3 convergent multi-donor class(*56*), while CR9114 is clustered with CR6261 and 315-02-1H01, which all shares V_H_1-69. The latter case is particularly noteworthy as CR9114 has much broader binding capacity than the other two, yet clustered with the stereotypical group 1-specific V_H_1-69 antibodies CR6261 and 315-02-1H01 (Figure 5).

**Fig. 5.**
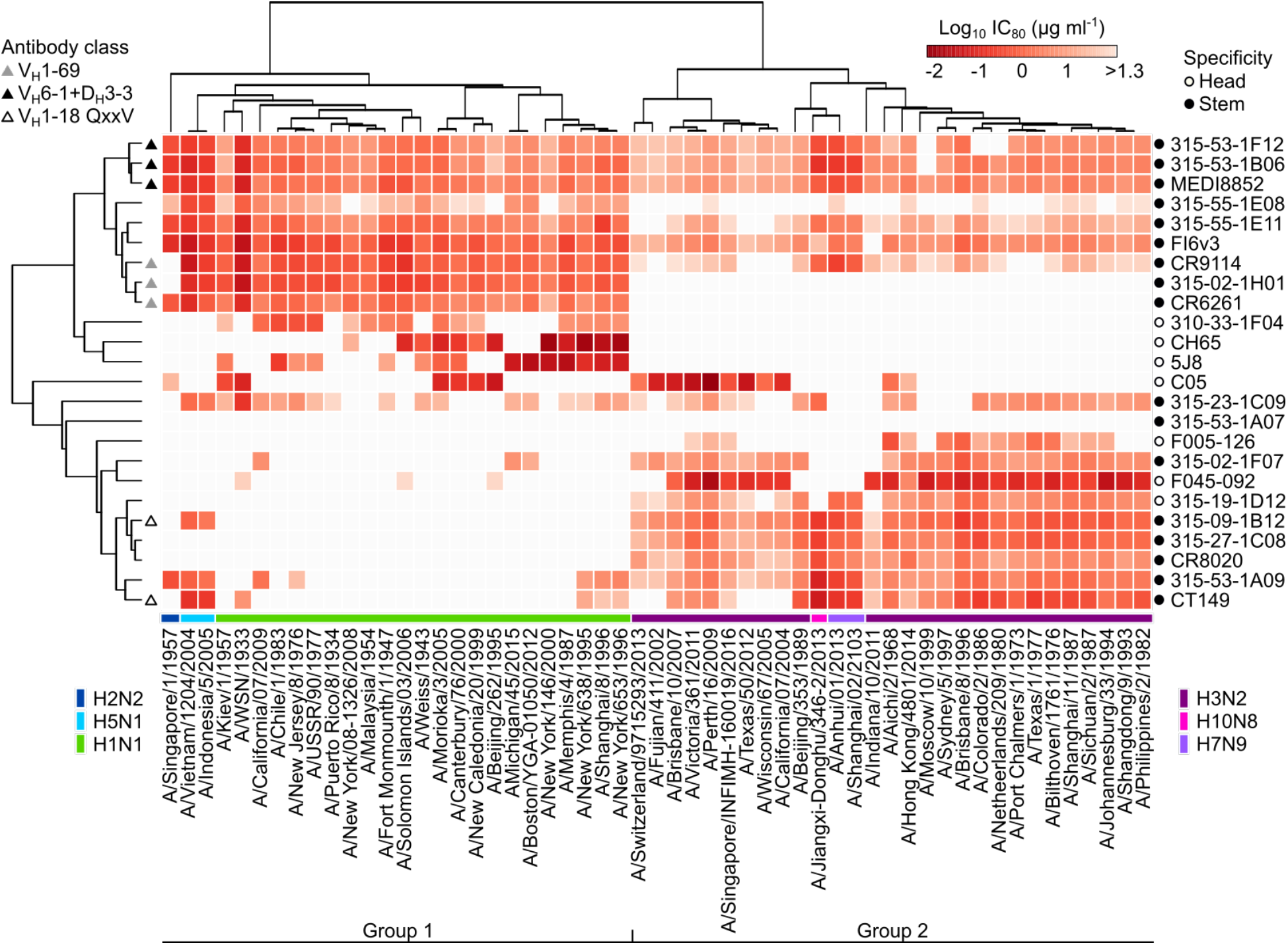
In-depth neutralization profiling of 24 monoclonal antibodies. Heatmap of neutralization titers (IC_80_ μg/ml) against a panel of 55 R3 influenza A viruses. Virus subtypes are color-coded and indicated on the bottom of the heatmap. Specificity of monoclonal antibodies is indicated on the right of the heatmap. Convergent classes of bnAbs with specific gene signatures are indicated on the left of the heatmap. Heatmap and unsupervised clustering was made with the ClustVis server (https://biit.cs.ut.ee/clustvis/).

Deep characterization of neutralizing profiles also predicts the developmental pathway for each bnAb. The convergent multi-donor bnAbs in V_H_1-18 QXXV class neutralize many group 2 viruses while having very limited breadth against group 1 viruses (Figure 5). This confirms previous findings in which the unmutated common ancestor (UCA) of this class of bnAbs engages only group 2 HAs and acquires group 1 reactivity through somatic hypermutation(*56*). Conversely, the V_H_6-1+D_H_3-3 class bnAbs possess higher neutralization potency against group 1 viruses than group 2 viruses (Figure 5), and this is consistent with the preferential engagement of group 1 HAs to the UCAs of this class(*56*). This deep neutralization profiling dataset also allows us to generate neutralization breadth-potency curves at relatively high resolution (Figure 6). Previous studies used a dataset generated by pseudotyped lentiviral neutralization assays with 15-17 selected HA-NA sequences(*40, 56*), and provided a limited understanding of neutralization breadth both because of the relatively small number of strains and the hypersensitivity of the pseudotyped lentivirus assay format. By determining the neutralization profiles using our comprehensive R3 virus panel, we found two antibodies (MEDI8852 and FI6v3) that were capable of neutralizing all 55 viruses (Figure 6). The other two V_H_6-1+D_H_3-3 class mAbs, 315-53-1F12 and 315-53-1B06, neutralized 52 (94.5%) and 54 (98.2%) of 55 viruses, respectively. Although CR9114 neutralized 48 out of 55 viruses (87.3%) its breadth-potency plot demonstrated a biphasic curve, highlighting the preferential neutralization of group 1 viruses (the first phase) and lower potency neutralization of group 2 viruses (the second phase) by this antibody (Figure 6). The receptor-binding site-directed antibody C05 showed limited neutralization breadth (32.7%) yet was extremely potent (Figure 6).

**Fig. 6.**
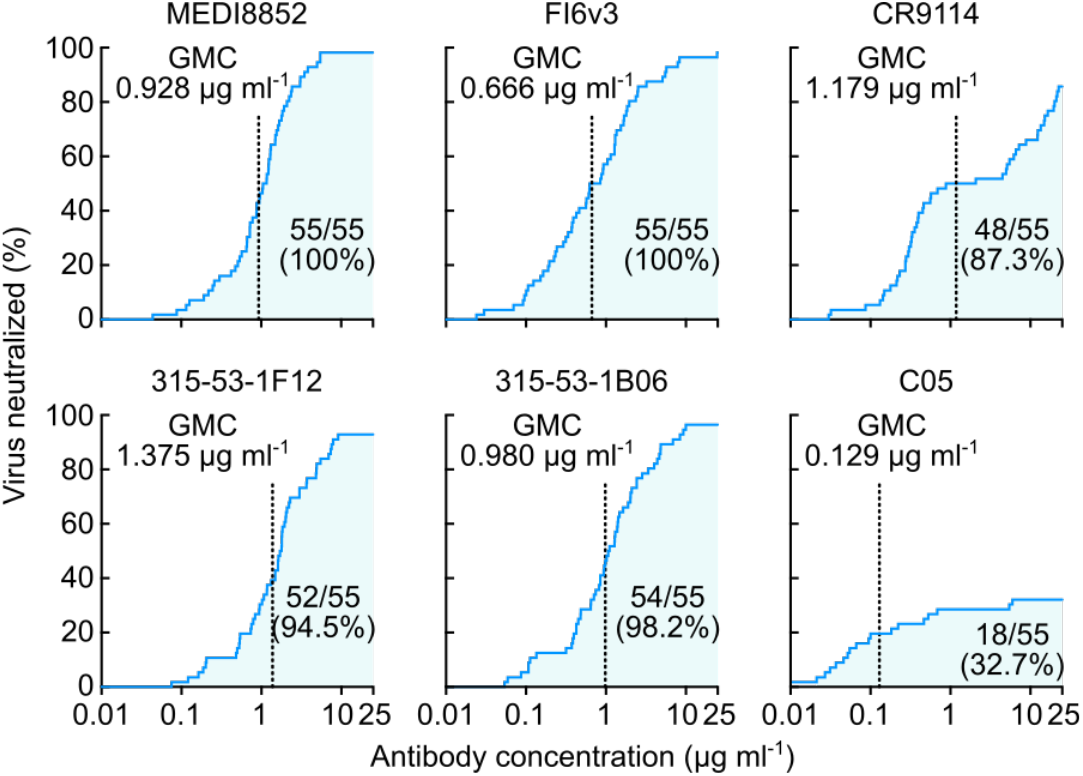
Neutralization breadth-potency analysis of broadly cross-reactive stem-directed antibodies and highly potent receptor-binding antibody. Neutralization titers (IC_80_ μg/ml) against a panel of 55 R3 viruses were used to generate the breadth-potency plot for each antibody. Geometric mean IC_80_ concentration (GMC) of each antibody was calculated only for the viruses neutralized by given antibody and indicated as a vertical dashed line on each graph. Shaded areas correspond to the number of viruses neutralized by given antibody. Both the number of viruses neutralized out of 55 R3 viruses and neutralization coverage (%) are indicated on each graph.

Overall, the neutralization profiling performed with the reporter influenza virus assay provides less biased breadth and potency information than the pseudotyped lentivirus assay or highly variable traditional MN assay. High resolution information may also help predict the class and the origin of antibodies with particular neutralization signatures.

## Discussion

Current influenza vaccines are imperfect and there is a large room to improve their efficacy, consistency, and breadth. Historically, efficacy has been associated with serum HAI activity(*57, 58*) and the HAI assay was developed as a surrogate for virus neutralization(*59, 60*). To improve current vaccines and to develop new vaccine concepts that would have broader and more durable efficacy, accurate measurement of neutralizing activity across a broad range of influenza viruses is critical. The HAI assay is a surrogate only for neutralization directing at the RBS on HA, therefore, neutralization targeting the HA stem or other relatively conserved sites on HA and NA will not be correlate with HAI. In addition, some new drifted seasonal strains of influenza and subtypes with pandemic potential do not always have predictable hemagglutination properties. Therefore, to encompass the variety of HA strains and the antibodies induced by diverse vaccine approaches, emphasis on a reproducible and authentic measurement of virus neutralizing activity is needed.

Unlike the HAI assay which measures the antibody’s ability to block receptor-binding of viral HA, plaque reduction(*61*) and MN assays(*62*) can capture neutralizing antibodies that block any virus replication step, including attachment, internalization, pH-induced conformational change of HA, membrane fusion, and virus egress. Despite efforts initiated by WHO, execution of influenza neutralization assays is still not standardized, leading to significant variability among results reported by different laboratories(*14, 62*). Development of a large panel of representative influenza viruses and significant improvement of assay throughput and safety are highly anticipated in the field to enable deep characterization and systematic comparisons of influenza virus neutralizing antibody responses. The virus panel combined with serum and monoclonal antibody standards would allow normalization across assay platforms and vaccine designs and facilitate the selection of universal influenza vaccine approaches to advance. In the present study, we address these outstanding needs by building an engineered influenza reporter virus-based neutralization assay coupled with high-throughput image-based readout in a biosafety level 2 setting. Our panel consists of 55 reporter viruses capturing almost the entire antigenic evolution of human H1N1 and H3N2 subtypes as well as historical human H2N2, and three other subtypes circulating at the human-animal interface.

Using the tdKatushka2 fluorescent reporter allowed us to measure virus replication in live cells without additional signal amplifications unlike ELISA- or hemagglutination-based readout. Image-based readout was chosen to further facilitated fast and precise data acquisition and analysis. In contrast to ELISA-based assays, in which the signal is measured for the entire well after a series of signal amplification steps, image-based detection provides higher signal-to-noise ratio, dynamic range, and precision by directly counting individual infected cells. The R3 virus neutralization assay can be readily automated to minimize hands-on time and human errors. Automation would facilitate standardization and increase reproducibility of the assay. It is also worth noting that the fluorescent image-based readout makes the assay more cost-effective compared with other assays such as ELISA- and luciferase-based assays.

As we continue to expand our collection of viruses and neutralizing antibodies, we will be able to perform neutralization fingerprint analysis(*63*) to provide a better understanding of the relationships between fine epitope specificity and neutralization breadth and potency, and allow computational predictions of the epitope-specific contributions of polyclonal serum antibodies to overall neutralizing activity. This type of analysis will foster the immune monitoring of antibody responses elicited by universal influenza vaccine candidates, particularly those targeting the conserved HA stem supersite(*64–66*).

As noted, previous reports have described the potential utility of R3ΔPB1(*23*) and R3ΔHA(*24, 25*) influenza viruses in neutralization assays. However, these efforts did not lead to the development of standardized assays due to a lack of comprehensive validation. We found that the antigenicity and neutralization sensitivity of R3ΔPB1 is similar to that of authentic influenza viruses, while R3ΔHA viruses appeared to be slightly more sensitive to neutralization than R3ΔPB1. This is likely due to less efficient HA incorporation into R3ΔHA viruses(*24, 25*). Although we noticed that the neutralization sensitivity of R4ΔPB1 closely matched that of R3ΔPB1 implying their potential superiority for building viruses that are incapable of reassortment, additional optimization of R4ΔPB1 virus rescue and propagation is necessary before expanding their practical applications.

Recent attempted improvements of the WHO-recommended ELISA-based MN assay protocol(*67–69*) underscore the need for the development of more reliable assays. Although the use of R3 viruses offers further advancements in the assay throughput, safety, and precision, this approach has its own limitations. Since the reporter segment is not required for viral replication, it is inevitable that after extensive virus passage, the reporter segment may accumulate mutations made by the viral error-prone viral RNA polymerase, which in turn, will diminish reporter gene activity and/or expression. However, it is possible to prepare virus stocks relatively quickly with five or less passages, and we confirm that these stocks retain an active reporter gene expression. Additionally, genomic stability of R3 influenza viruses can be improved by utilizing the variant PB1 polymerase with lower error-rates as reported recently(*70*) and designing R3 viruses capable of replicating only when the engineered genomic segment is functional, such as inducible gene-expression systems. Alternatively, plaque purification or rederiving virus strains may be necessary periodically. By describing our methods and depositing sequences required to generate R3 viruses, we provide the necessary information to implement this technology for the strains reported here or future emerging viruses. Nevertheless, the R3 influenza virus neutralization assay reported here substantially improves existing MN assays and fosters rapid discovery and characterization of influenza bnAbs as well as vaccine-elicited serum antibody responses to support the development of universal influenza vaccines.

## Acknowledgments

We thank Richard Webby (St. Jude Research Hospital) for influenza reverse genetics plasmids and Jesse Bloom (Fred Hutchinson Cancer Research Center) for HEK-293 cells expressing PB1 of A/WSN/1933.

## Author contributions

Conceptualization: A.C., M.K., B.S.G.; Methodology: A.C.; Formal Analysis: A.C., M.K.; Investigation: A.C., R.A.G., B.E.F., S.F.A., L.H., T.S., Y.T.; Writing – Original Draft: A.C., M.K.; Writing – Review & Editing: A.C., S.F.A., J. R.M., B.S.G., M.K.; Supervision: A.B.M., J.R.M., B.S.G., M.K.; Project Administration: M.C.C.; Funding Acquisition: J.R.M., B.S.G.

## Data and materials availability

Influenza reverse genetics plasmids were obtained from St. Jude Research Hospital thorough an MTA. All sequences corresponding to the influenza HA, NA and reporter segments used in the present study have been deposited to NCBI Genbank under accession numbers xxxxxxxx–xxxxxxxx.

## Supplementary Materials

Materials and Methods

Fig. S1 – S4.

References (71 –73)

