## Supplementary materials for "A comprehensive influenza reporter virus panel for high-throughput deep profiling of neutralizing antibodies"

**The PDF file includes:**

Materials and Methods

Fig. S1. Sequence of PB1 segment of R3ΔPB1 influenza viruses.

Fig. S2. Growth kinetics of R3ΔPB1 virus in cells with or without PB1 expression.

Fig. S3. Neutralization assay using R3ΔPB1 influenza virus.

Fig. S4. Generation of rewired replication-restricted reporter (R4) ΔPB1 influenza virus.

### Materials and Methods

*Plasmids*

To prepare influenza reverse genetics plasmids for rescue of influenza A H1N1 or H3N2 viruses described in this study, HA and NA coding sequences were retrieved from Genbank; noncoding regions of A/WSN/1933 for H1N1 or A/Netherlands/009/2010 for H3N2 viruses were added at both ends. Full-length HA and NA sequences were cloned into a dual promoter influenza reverse genetics plasmid previously described (Supplemental reference 1). To rescue influenza viruses, dual promoter plasmids encoding internal genes of A/WSN/1933 (pHW181-PB2, pHW182-PB1, pHW183-PA, pHW185-NP, pHW187-M, pHW188-NS) were used(*71*). To prepare PB1 reporter segment used to rescue replication-restricted reporter viruses, the sequence containing the PB1 genome packaging signals of A/WSN/1933(*27*) and mKate2 or tdKatushka2 reporter coding region(*28*) (Addgene Cat No. 56049) was synthesized and cloned using BsmBI (New England Biolabs) restriction sites into the dual promoter influenza reverse genetics plasmid. Similarly, HA reporter segment was synthesized with HA genome packaging signals of A/Puerto Rico/1934(*24*) flanking the tdKatushka2 reporter sequence and cloned into the dual promoter influenza reverse genetics plasmid using BsmBI restriction sites. To prepare the influenza reverse genetics plasmid using chicken beta-actin CAG pol-II promoter (Addgene Cat No. 41583), an insert comprising human pol-I promoter and mouse pol-I terminator sequences in negative orientation flanking two BsmBI restriction sites was cloned using KpnI and XhoI restriction sites. Then, full-length of influenza genes of high-yield A/Puerto Rico/8/1934 were cloned into BsmBI restriction sites(*72*). To prepare plasmids for stable cell line development, sequences of Streptomyces puromycin N-acetyl-transferase (PAC), which confers resistance to puromycin, followed by self-cleaving peptide of *Thosea asigna* virus 2A (T2A) and coding region of influenza A PB1 or HA genes were synthesized and cloned using KpnI and XhoI restriction sites. All plasmids were confirmed by Sanger sequencing.

*Cells*

To propagate influenza viruses, MDCK-SIAT1 cells (Sigma) were used. Cells were maintained with complete media comprising Dulbecco's modified Eagle's medium high glucose (DMEM; Thermo Fisher Scientific) supplemented with 10% (v/v) heat-inactivated fetal bovine serum (Gemini Bio-Products), 100 units/ml penicillin (Thermo Fisher Scientific), 100 µg/ml streptomycin (Thermo Fisher Scientific) and geneticin (1 mg/ml) (Thermo Fisher Scientific). To develop constitutively PB1- or HA-expressing MDCK-SIAT1 cells, one plasmid encoding both puromycin resistance and influenza genes was transfected into MDCK-SIAT1 cells using Lipofectamine 2000. Two days post transfection, cells were transferred from 6 well plates into 10-cm dishes containing DMEM media with 10% bovine serum (Gemini Bio-Products), penicillin, streptomycin, geneticin (1 mg/ml) (Thermo Fisher Scientific) and puromycin (0.25 μg/ml) (Thermo Fisher Scientific) for selection. Medium was changed every 48 hours. Clonal selection was performed using 8 or 10 mm cloning cylinders (Fisher Sciences) about two weeks after the transfection. Clonal cell lines were screened using a reporter virus prepared on a polyclonal cell line. To rescue influenza viruses, FlpIn HEK-293 cells (Thermo Fisher Scientific) were transfected as described below.

*Reverse genetics of influenza viruses*

For molecular clone influenza viruses, an eight-plasmid approach in which each influenza segment is inserted between pol-II (positive orientation) and pol-I (negative orientation) promoters was used to rescue parental molecular clone viruses. Briefly, FlpIn HEK-293 (Thermo Fisher Scientific) cells were transfected with dual promoter plasmids encoding each influenza segment of A/WSN/1933 (obtained from St. Jude Children's Research Hospital) and a CMV-driven plasmid expressing human transmembrane serine protease 2 (hTMPRSS2). Transfection was performed using Lipofectamine 3000 in 6-well plates coated with D-lysine. Three days post infection, TPCK-trypsin (0.5 μg/ml) was added to the transfected cells for 2-4 hours. Supernatant was harvested, cleared by centrifugation (200 × *g*, 10 min) and used for propagation in MDCK-SIAT1 cells by limiting dilution. For virus propagation, the inoculum was prepared in virus growth medium comprising of OptiMEM (Thermo Fisher Scientific) supplemented with TPCK-trypsin at 1 μg/ml (Sigma). MDCK-SIAT1 flasks containing cells at ~80% confluence were washed twice with phosphate-buffered saline (PBS) and incubated with the inoculum at 37°C for virus adsorption. After 1 h, the inoculum was removed, cells were washed with PBS and virus growth media was added to the cells. The infected cells were incubated at 37°C in a humidified 5% CO2 atmosphere. After two days, supernatant was harvested when CPE reached 40-60%, cleared by centrifugation (200 × *g*, 10 min), aliquoted and stored at –80°C. For influenza HA and NA sequencing, viral RNA was extracted with Qiagen RNeasy extraction kit (Qiagen). Influenza viral RNA was amplified using the Qiagen One-Step RT-PCR kit (Qiagen) and HA and NA specific primers tagged with M13 sequences. Each amplicon was sequenced with M13 primers in both directions. Primer sequences are available upon request. Sequences were analyzed using Sequencer 5.4 (Gene Codes).

For replication-restricted reporter ΔPB1 influenza viruses, FlpIn HEK-293 (Thermo Fisher Scientific) cells were transfected with eight dual promoter influenza reverse genetics plasmids (the PB1 reporter segment plasmid replaced the plasmid encoding PB1 of A/WSN/1933) together with pol-II-driven expression plasmid encoding PB1 of A/WSN/1933 and hTMPRSS2. Rescued viruses were propagated in MDCK-SIAT1 cells constitutively expressing PB1 of A/WSN/1933 in the presence of TPCK-trypsin (1 μg/mL, Sigma). Virus stocks were stored at –80$^{o}$C.

For replication-restricted rewired ∆PB1 influenza viruses FlpIn HEK-293 (Thermo Fisher Scientific) cells were transfected with eight dual promoter influenza reverse genetics plasmids (PB1 segment comprises PB1 genome packaging signals flanking the coding region of an HA segment in which the HA genome packaging signals have been destroyed by introducing silent mutations, while the HA segment of R4∆PB1 contains the reporter gene flanked by intact HA genomic packaging signals) together with pol-II-driven expression plasmid encoding PB1 of A/WSN/1933 and hTMPRSS2. Rescued viruses wre propagated as described above.

For replication-restricted reporter ∆HA influenza genes, HEK-293 cells were transfected with eighth influenza reverse genetics plasmids encoding sequences of high-yield A/Puerto Rico/8/1936 (HA segment was replaced with HA reporter segment described above), pol-II-driven HA and hTMPRSS2 expressing plasmids. Viruses were propagated in MDCK-SIAT1 cells constitutively expressing influenza HA gene in the presence of TPCK-trypsin (1µg/mL, Sigma). Virus stocks were stored at –80$^{o}$C

*Phylogenetic and evolution-based conservation analyses of influenza HA*

Nucleotide sequences of mature H1 HA (N=25) and H3 HA (N=26) proteins were aligned using Muscle algorithm found in Bioedit v7.2.5. Phylogenetic trees were generated using neighbor-joined methods and Kimura 2-parameter substitution model as implemented in MEGA v10. Evolution-based conservation analyses of amino acid residues in the extracellular region of H1 and H3 HA proteins was done using Consurf (http://consurf.tau.ac.il) and visualized on the atomic structures of HA proteins of A/California/07/2009 (PDB ID: 3LZG) and A/Victoria/361/2011 (PDB ID: 4WE8). To evaluate the conservation of each amino acid of H3 HA proteins isolated from influenza viruses circulating in humans, full-length H3 HA sequences of human H3N2 viruses were obtained from the GISAID database (http://platform.gisaid.org). Alignment of nucleotide sequences was performed using MAFFT v7 server-based algorithm using default settings (https://mafft.cbrc.jp/alignment/server/large.html). After removal of redundant sequences and sequences with gaps or degenerate nucleotide bases, we obtained a dataset of 16,893 unique sequences. Due to the large size of this sequence data set, we clustered sequences (CD-HIT Suite at http://weizhong-lab.ucsd.edu/cdhit-web-server/cgi-bin/index.cgi?cmd=cd-hit-est) with identity higher than 99.6% identity and choose a representative sequence from each cluster. The final dataset used to estimate evolution-based conservation of amino acid residues in human H3 HA proteins had 3,408 sequences. Similarly, an alignment of 1,222 H1 HA of H1N1 viruses circulating in humans between 1918 and 2019 sequences was used to evaluate the conservation of each amino acid of this protein.

*Virus titration and neutralization*

For molecular clones, ELISA-based influenza microneutralization assay was performed following WHO-recommended protocol. Briefly, influenza viruses were titrated in MDCK-SIAT1 cells plated in 96-well plates at 50,000 cells/ml 24 h before the infection. Virus dilutions were prepared using OPTIMEM (Thermo Fisher) supplemented with TPCK trypsin (Sigma) at a 1µg/mL final concentration. Infected cells were incubated at 37°C and 5% CO2 humidified atmosphere. After 18 h, cells were fixed 80% cold acetone and air dried. Virus replication was detected by ELISA with biotin-conjugated antibodies to influenza virus nucleoprotein (MAB8257B and MAB8258B, EMD Millipore) and was visualized with HRP-conjugated streptavidin and SureBlue TMB Microwell Peroxidase Substrate (KPL). Absorbance was read at 450 nm (A450) and 650 nm (A650) with the SpectraMax Paradigm microplate reader (Molecular Devices). The A650 was used to subtract plate background. $TCID_{50}$ titer was calculated using Reed-Much algorithm. Neutralization assays were performed using 100-200 $TCID_{50}$ units of virus and 4-fold antibody dilutions made in OptiMEM (Thermo-Fisher). TPCK-trypsin was added to a final concentration of 1 µg/mL. Virus and antibody were mixed in equal volumes and incubated 1 h at 37°C prior to adding to substrate MDCK-SIAT1 cells. Control wells of virus alone (VC) and diluent alone (CC) were included on each plate. Fifty microliters of antibody/virus mixture were then added to wells of pre-washed cells in duplicate and the plates were incubated for 18 h at 37°C and 5% $CO_2$ humidified atmosphere. Infected cells were detected as described above. The percent neutralization was calculated by constraining the VC control as 0% and the CC control as 100% and plotted against serum/antibody concentration. A curve fit was generated by a four-parameter nonlinear fit model in Prism (GraphPad). The 80% ($IC_{80}$) inhibitory concentrations were obtained from the curve fit for each serum sample or antibody respectively.

Titer of replication-restricted reporter ∆PB1 or ∆HA viruses was measured in PB1-expressing MDCK-SIAT1 cells plated in 96-well black plates with transparent bottom (Greiner) at 18 hours post infection and counting fluorescent-foci using Celigo Image-Cytometer (Nexcelom) with customizied red channel to enhance detection of mKate2/TdKatushka2 reporter (EX 540/80 nm, DIC 593 nm and EM 593/LP nm). The Celigo operation and analysis software consisted of five major steps START, SCAN, ANALYZE, GATE and RESULTS, where the user can enter setup scan and analysis parameters. Target 1 protocol was used to detect and count fluorescent foci. Titer was expressed as fluorescent foci per ml. For each neutralization reaction, the virus dilution that resulted in cca. 1000 (500-4,000) fluorescent foci per well at 18 h post infection was used. Neutralization assays using R3 viruses performed to compare the fluorescence- and ELISA-based assays were done in 96-well black plates. Cells were plated 24 h before the experiment. Neutralization reaction was done as described above. R3 influenza neutralization assay was optimized to be performed in 384-well plate format. PB1-expressing MDCK-SIAT1 cells were washed twice with PBS, re-suspended in OPTIMEM and plated 2 h before the assay in 384-well plates at 150,000 cells/ml with each well containing twenty microliters. Twenty-five microliters of each neutralization mix consisted of 2 μg/mL TPCK trypsin and equal parts of virus and 4-fold serial dilutions of monoclonal antibodies were transferred to wells in quadruplicate. Control wells of virus alone (VC) and diluent alone (CC) were included on each plate. Fluorescent foci were counted at 18-24 h post infection using an image-based plate reader. Neutralization titers were calculated as using Prism (GraphPad) as described above.

*Antibody preparation*

Sequences of immunoglobulin heavy and light chains were synthesized and cloned into human IgG1 as previously described(*48,73*). The expression vectors were transiently transfected into Expi293F (Thermo Fisher Scientific) using ExpiFectamine 293 transfection reagents (Thermo Fisher Scientific). Monoclonal antibodies were purified using sepharose Protein-A or G (Pierce) following manufacturer's instructions.

*Negative stain EM*

Virus preparations were mixed at a 1:1 ratio with fixative containing 4% glutaraldehyde and 0.2 M cacodylate buffer, pH 7. A drop of the fixed sample was placed on a carbon-coated, glow-discharged copper grid for about 30 s. The drop was then removed using filter paper, and the grid was washed with three drops of buffer containing 10 mM HEPES, pH 7, and 150 mM NaCl, followed by negative staining with 0.75% uranyl formate. Imaging was performed using a ThermoFisher Talos F200C electron microscope operated at 200 kV and equipped with a Ceta camera.

*Statistical Significance*

All statistical analysis was performed using Prism Graphpad software. Specific tests to determine statistical significance used are indicated in the methods and corresponding figure legends. Probability values less than 0.05 were considered statistically significant.

*5'-*
*agcgaaagcaggcaaaccatttga*TtggTtgtcaatccgactttacttttcttaaaagtgccagcacaaaTtgctataagcacaactttcccttatactggagaccctccttacagccTtgggacaggaacaggatac
accTtgggtaccATGGTGGGTGAGGATAGCGTGCTGATCACCGAGAACATGCACATGAAACTGTACATGGAGGGCACCGTGAACGACCACCACTTCAA
GTGCACATCCGAGGGCGAAGGCAAGCCCTACGAGGGCACCCAGACCATGAAGATCAAGGTGGTCGAGGGCGGCCCTCTCCCCTTCGCCTTCGA
CATCCTGGCTACCAGCTTCATGTACGGCAGCAAAACCTTTATCAACCACACCCAGGGCATCCCCGACTTCTTTAAGCAGTCCTTCCCTGAGGGCTT
CACATGGGAGAGGATCACCACATACGAAGACGGGGGCGTGCTGACCGCTACCCAGGACACCAGCCTCCAGAACGGCTGCCTCATCTACAACGTC
AAGATCAACGGGGTGAACTTCCCATCCAACGGCCCTGTGATGCAGAAGAAAACACTCGGCTGGGAGGCCAGCACCGAGATGCTGTACCCCGCTG
ACAGCGGCCTGAGAGGCCATGCCCAGATGGCCCTGAAGCTCGTGGGCGGGGGCTACCTGCACTGCTCCCTCAAGACCACATACAGATCCAAGAA
ACCCGCTAAGAACCTCAAGATGCCCGGCTTCTACTTCGTGGACAGGAGACTGGAAAGAATCAAGGAGGCCGACAAAGAGACCTACGTCGAGCAG
CACGAGATGGCTGTGGCCAGGTACTGCGACCTGCCTAGCAAACTGGGGCACAGCGGGCATGGCACCGGCAGCACCGGCAGCGGCAGCTCCGG
TACtGCCTCCTCCGAGGACAACAACATGGCCATGGTGGGTGAGGATAGCGTGCTGATCACCGAGAACATGCACATGAAACTGTACATGGAGGGCAC
CGTGAACGACCACCACTTCAAGTGCACATCCGAGGGCGAAGGCAAGCCCTACGAGGGCACCCAGACCATGAAGATCAAGGTGGTCGAGGGCGG
CCCTCTCCCCTTCGCCTTCGACATCCTGGCTACCAGCTTCATGTACGGCAGCAAAACCTTTATCAACCACACCCAGGGCATCCCCGACTTCTTTAA
GCAGTCCTTCCCTGAGGGCTTCACATGGGAGAGGATCACCACATACGAAGACGGGGGCGTGCTGACCGCTACCCAGGACACCAGCCTCCAGAAC
GGCTGCCTCATCTACAACGTCAAGATCAACGGGGTGAACTTCCCATCCAACGGCCCTGTGATGCAGAAGAAAACACTCGGCTGGGAGGCCAGCA
CCGAGATGCTGTACCCCGCTGACAGCGGCCTGAGAGGCCATGCCCAGATGGCCCTGAAGCTCGTGGGCGGGGGCTACCTGCACTGCTCCCTCA
AGACCACATACAGATCCAAGAAACCCGCTAAGAACCTCAAGATGCCCGGCTTCTACTTCGTGGACAGGAGACTGGAAAGAATCAAGGAGGCCGAC
AAAGAGACCTACGTCGAGCAGCACGAGATGGCTGTGGCCAGGTACTGCGACCTGCCTAGCAAACTGGGGCACAGCTCCGGACTCAGATCTTGAct
cgaggtttccagagcccgaattgatgcacgaattgatttcgaatctggaaggataaagaaagaggagttcactgagatcatgaagatctgttccaccattgaagagctcagacggcaaaaatag*tgaatttagcttgtc*
*cttcatgaaaaaatgccttgtttctact-3'*

Italics: NCR
Red: TdKatushka2 ORF
Underlined: restriction sites (KpnI/XhoI)
Capital letters (black): mutated ATG in the coding packaging region

**Fig. S1.** Sequence of PB1 segment of R3ΔPB1 influenza viruses. Sequences corresponding to tdKatushka2 reporter is in red, non-coding regions are italicized, coding region of the packaging signals are in black, mutated ATG start codons in the 5' PB1 packaging signal are indicated with capital letters, restriction sites are underlined.

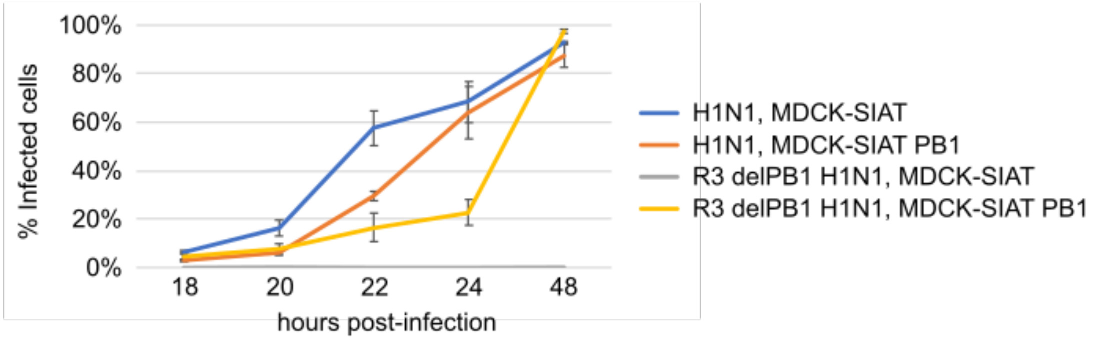


**Fig. S2**. Growth kinetics of R3ΔPB1 virus in cells with or without PB1 expression. MDCK-SIAT1 and PB1-expressing MDCK-SIAT1 were infected with R3ΔPB1 or the parental A/Michigan/45/2015 (H1N1) viruses. Infected cells fixed with 4% paraformaldehyde (Electron Microscopy Sciences) and permeabilized with 0.01% Triton-X (Sigma) were detected by ELISA with biotin-conjugated antibodies to influenza virus nucleoprotein (MAB8257B and MAB8258B, EMD Millipore) and imaged with streptavidin coupled with Alexa 488 (Theromo Fisher). DAPI (300 nM) (Thermo Fisher) was used to label all cells. Stained cell populations were counted automatically using Celigo.

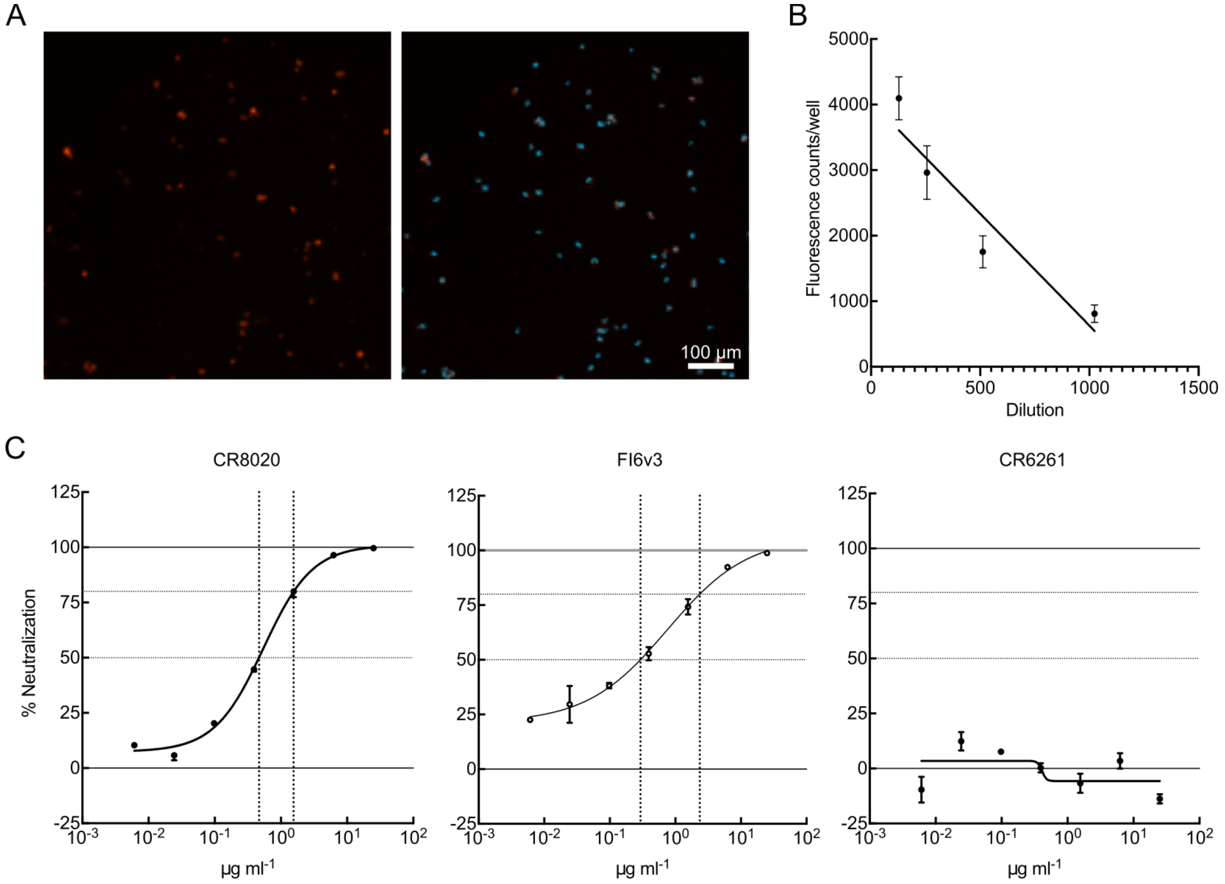


**Fig. S3**. Neutralization assay using R3ΔPB1 influenza virus. (A) R3ΔPB1 A/Singapore/INFIMH-16-0019/2016 (H3N2) virus-infected MDCK-SIAT1-PB1 cells at 18 hours post infection. Images representing single wells of 384-well plate are acquired using Target 1 protocol of Celigo. Fluorescent foci (left panel: red foci) are identified and counted automatically (right panel: foci with blue contour) using manufacturer software. (B) Titration of R3ΔPB1 A/Singapore/INFIMH-16-0019/2016 (H3N2) virus. Linear range of fluorescence readings against virus dilutions is shown ($R^2 = 0.8519$). (C) Representative neutralization profiles of three mAbs against R3ΔPB1 A/Singapore/INFIMH-16-0019/2016 (H3N2) virus. Normalized readings with standard deviations and fitted curves are shown for each mAb. The 50% ($IC_{50}$) and 80% ($IC_{80}$) inhibitory concentrations are obtained for each antibody from the fitted curve and shown with dotted lines.

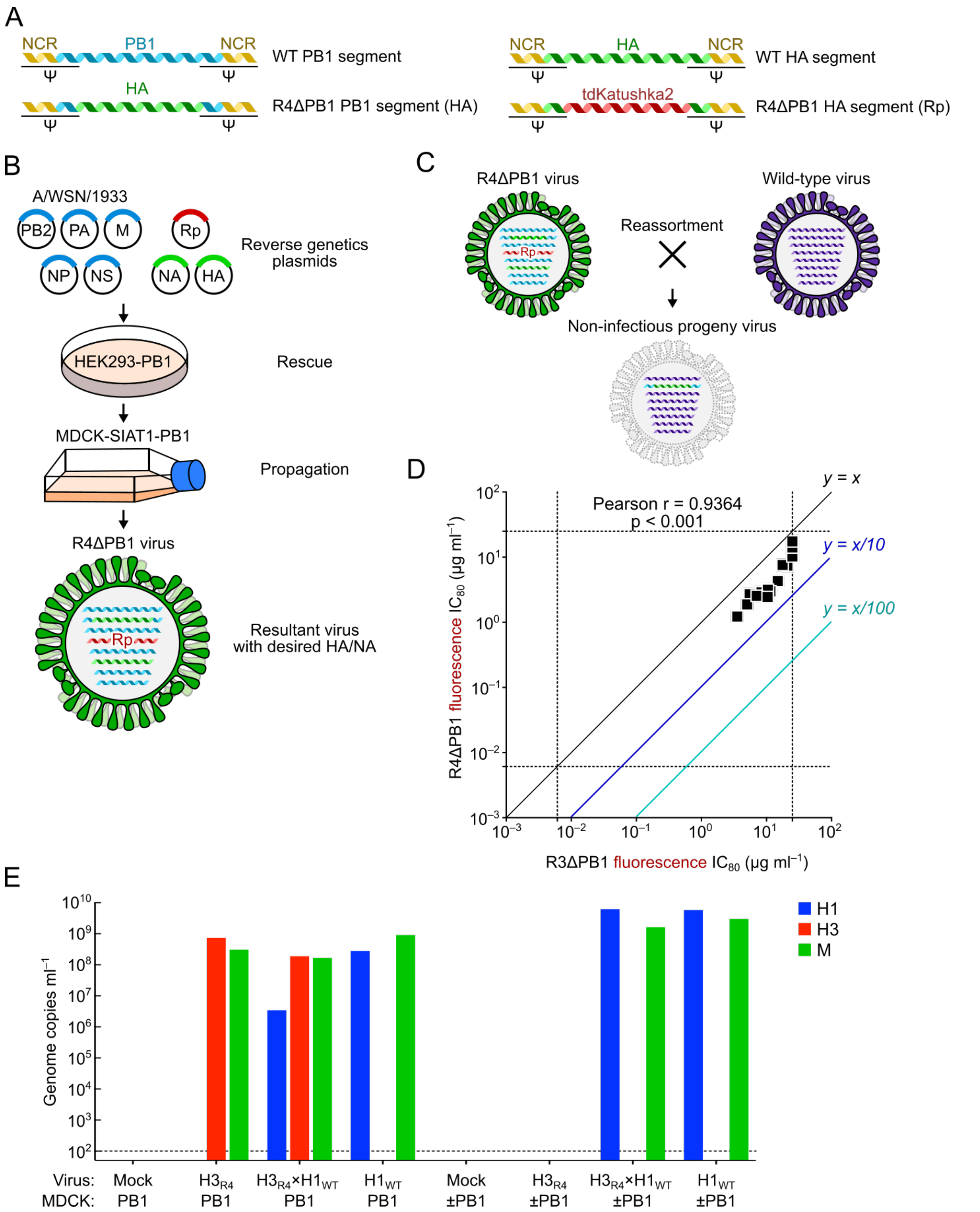


**Fig. S4.** Generation of rewired replication-restricted reporter (R4) ΔPB1 influenza virus. (A) Design of modified PB1 and HA segments used for R4ΔPB1 virus. PB1 segment contains PB1 packaging signals and HA ORF with mutated packaging signals. HA segment contains the HA packaging signals flanking the reporter ORF. (B) R4ΔPB1 virus rescue and propagation. (C) Non-viable reassortment between R4ΔPB1 and wild-type influenza viruses. Reassortant virus carrying the engineered PB1 segment encoding HA ORF and wild-type HA segment results in replication-deficient virus due to lack of PB1 gene. (D) Correlation between neutralization titers of 24 mAbs against R3ΔPB1 and R4ΔPB1 viruses (A/Switzerland/9715293/2013). Each dot indicates titers ($IC_{80}$ μg $ml^{-1}$) of a single mAb against R4ΔPB1 (y-axis) and R3ΔPB1 (x-axis) determined by fluorescent readout. (E) qRT-PCR analysis of influenza viruses to detect reassortment event. Forced reassortment experiment between R4ΔPB1 H3N2 (A/Switzerland/9715293/2013, $H3_{R4}$) and H1N1 (A/Solomon Islands/03/2006, $H1_{WT}$) was performed in MDCK-SIAT1 expressing PB1 cells followed by propagation in parental MDCK-SIAT1 cells. MDCK SIAT1 expressing PB1 were infected with $H3_{R4}$, $H1_{WT}$ or 1:1 mixture of the two viruses. Due to the low titer of $H3_{R4}$ (6,270 $TCID_{50}$ $ml^{-1}$), the first passage was done at MOI of 0.2. Supernatants were harvested 48 hours post-infection. Viruses were initially passaged 3 times on MDCK-SIAT1 expressing PB1 to maximize the chances of reassortment events between $H3_{R4}$ and $H1_{WT}$ viruses, and then passaged 6 times on parental MDCK-SIAT1 cells (no PB1) to allow propagation of reassortant viruses (labeled as ±PB1). Viral RNA extracted from supernatants was analyzed by qRT-PCR with primers and probes designed for the detection of H1 HA (blue), H3 HA (red), and M (green) genes. Viral RNA standard of A/Puerto Rico/8/1934 (ATCC: VR-95PQ) and A/Hong Kong/8/1963 (H3N2) (ATCC: VR-544PQ), that have the genome copy number determined by droplet digital PCR were used as standard for qRT-PCR.
